# Core Circadian Clock Genes *Per1* and *Per2* regulate the Rhythm in Photoreceptor Outer Segment Phagocytosis

**DOI:** 10.1101/2020.12.23.424210

**Authors:** Nemanja Milićević, Ouafa Ait-Hmyed Hakkari, Udita Bagchi, Cristina Sandu, Aldo Jongejan, Perry D. Moerland, Jacoline B. ten Brink, David Hicks, Arthur A. Bergen, Marie-Paule Felder-Schmittbuhl

## Abstract

Retinal photoreceptors undergo daily renewal of their distal outer segments, a process indispensable for maintaining retinal health. Photoreceptor Outer Segment (POS) phagocytosis occurs as a daily peak, roughly about one hour after light onset. However, the underlying cellular and molecular mechanisms which initiate this process are still unknown. Here we show that, under constant darkness, mice deficient for core circadian clock genes (*Per1* and *Per2*), lack a daily peak in POS phagocytosis. By qPCR analysis we found that core clock genes were rhythmic over 24h in both WT and *Per1, Per2* double mutant whole retinas. More precise transcriptomics analysis of laser capture microdissected WT photoreceptors revealed no differentially, expressed genes between time-points preceding and during the peak of POS phagocytosis. By contrast, we found that microdissected WT retinal pigment epithelium (RPE) had a number of genes that were differentially expressed at the peak phagocytic time-point compared to adjacent ones. We also found a number of differentially expressed genes in *Per1, Per2* double mutant RPE compared to WT ones at the peak phagocytic time-point. Finally, based on STRING analysis we found a group of interacting genes which potentially drive POS phagocytosis in the RPE. This potential pathway consists of genes such as: *Pacsin1*, *Syp*, *Camk2b* and *Camk2d* among others. Our findings indicate that *Per1* and *Per2* are necessary clock components for driving POS phagocytosis and suggest that this process is transcriptionally driven by the RPE.

**Declarations:** *Funding:* This project has been funded with support from the NeuroTime Erasmus+ grant (European Union), Rotterdamse Stichting Blindenbelangen (Netherlands), Nelly Reef fund (Netherlands), Stichting voor Ooglijders (Netherlands), Stichting tot Verbetering van het Lot der Blinden (Netherlands) and Retina France (France).

*Conflicts of interest/Competing interests:* The authors declare no competing interests.

*Availability of data and material:* Data supporting the conclusions of this article are included within the article and are available from the corresponding authors on reasonable request.

*Code availability:* The R code for analysis is available from the corresponding authors on reasonable request.

*Ethics approval:* All experimental procedures were performed in accordance with the Association for Research in Vision and Ophthalmology Statement on Use of Animals in Ophthalmic and Vision Research, as well as with the European Union Directive (2010/63/EU).

*Consent to participate:* Not applicable

*Consent for publication:* All authors read and approve of the contents of this manuscript.

*Author contributions:* N.M. performed experiments, analysis, prepared figures, wrote the manuscript and obtained funding. O.A.-H.H. performed experiments, data analysis, prepared figures and obtained funding. P.D.M. and A.J. performed bioinformatics analysis and edited the manuscript. U.B., J.B.t.B. and C.S. provided technical assistance, performed experiments, prepared figures and edited the manuscript. D.H., A.A.B. and M.-P.F.-S. conceptualized and directed the project, obtained funding, provided resources, performed analysis and edited the manuscript.

## Introduction

Light/dark transitions are one of the hallmarks of life on Earth. Living organisms adapt their behavior and physiology according to cyclic changes in environmental conditions. In mammals, these rhythmic adjustments in molecular and cellular physiology are enabled through a hierarchical network of oscillators, encompassing a “central clock” located in the suprachiasmatic nucleus (SCN) in the brain and peripheral oscillators [1]. The core molecular components generating these oscillations are comprised of interlocking transcriptional-translational feedback loops involving “clock” transcription factors such as PER1-2, CLOCK, BMAL1, CRY1-2, REV-ERBs and RORs [2]. These factors drive rhythmic expression of “clock-controlled genes” thereby enabling rhythmic adaptations in physiology.

The retina stands out as a peripheral oscillator as it lies in direct contact with the main environmental synchronizing stimulus – light [3]. This light-sensitive organ is composed of multiple layers of cells, all of which were shown to oscillate in a layer-specific manner and are strongly coupled [4]. Numerous aspects of retinal physiology and functions were shown to be rhythmic [5] such as melatonin release [6,7], rod-cone coupling [8,9], visual sensitivity [10,11] and photoreceptor disc shedding [12]. Of all retinal cells, circadian oscillations in photoreceptors have been most extensively studied (reviewed by [13]).

Retinal photoreceptors are specialized, light-sensitive neuronal cells. They are metabolically highly active cells in which homeostasis is tightly controlled. They consist of a cell body, a specialized synapse, inner and outer segments. Together with the adjacent retinal pigment epithelium (RPE), the POS contain the molecular machinery that sustains phototransduction. Excessive light exposure can damage these cells. A mechanism that prevents the accumulation of photo-oxidative compounds is rapid POS renewal [14]. This turnover involves several critical steps. At the proximal POS end, these steps include synthesis and intracellular transport of structural and functional proteins. At the distal end, POS fragments are shed and subsequently phagocytosed by the RPE. Impairment of phagocytosis was previously implicated in photoreceptor degeneration in both animal models [15] and humans [16]. Despite many studies devoted to the subject, the molecular mechanisms that control POS phagocytosis remain elusive [5,17,18]. Phagocytosis of POS was shown to be highly cyclic, taking place in rods as a daily peak occurring about one hour after light is turned on in both nocturnal and diurnal mammals [12,19,20]. This peak is maintained under constant darkness, implicating circadian control. However, little is known about the transcriptional events that occur prior and during the peak of POS phagocytosis.

In the present study, we tested the hypothesis that *Per1* and *Per2* are necessary clock components for initiating the phagocytosis of rod outer segment in mice. We investigated the transcriptional changes that occur in the RPE and photoreceptors prior and during the peak in POS phagocytosis. Finally, we proposed a potential pathway for initiating POS phagocytosis based on our transcriptomics data obtained from multiple time-points, purest possible microdissected sample material and phagocytically arrhythmic *Per1, Per2* mouse double knockout model.

## Methods

### Animals

Experiments were conducted using homozygote double mutant mice carrying the loss-of-function mutation of *mPer1* gene (*Per1^−/−^*; [21]) and mutation of the *mPer2* gene (*Per2^Brdm1^*, [22]; hereafter defined as *Per1^−/−^ Per2^Brdm1^* or KO). Intercrosses between heterozygous (C57BL/6/J × 129 SvEvBrd) F1 offspring gave rise to F2 homozygous mutants. Mutant and wild-type (WT) animals on this mixed background were used in this study, maintained as described in [23]. Mice were maintained in our animal facilities (Chronobiotron, UMS3415, Strasbourg, France) on a 12h light/12h dark (LD) cycle (300 lux during the light phase), with an ambient temperature of 22 ± 1 °C. The animals were given free access to food and water. In all experiments, control and mutant mice were age-matched. Only male mice were used for the RNAseq study, but both males and females were used for qPCR experiments and phagocytosis analysis. All experimental procedures were performed in accordance with the Association for Research in Vision and Ophthalmology Statement on Use of Animals in Ophthalmic and Vision Research, as well as with the European Union Directive (2010/63/EU). Age-matched WT and *Per1^−/−^Per2^Brdm1^* mice (6 weeks old) were sacrificed in constant darkness (dark/dark, DD) at time-points (expressed in circadian time (CT); CT0 – time when lights were on during LD conditions, CT12 – lights off in LD conditions) specific to each experiment. Sacrifice was performed under complete darkness by using night-vision goggles ATN NVG-7 (American Technologies Network Corp., San Francisco, CA, USA) and eye sampling was done under dim red light (< 5 lux). Animals were anesthetized by CO2 inhalation and subsequently killed by cervical dislocation.

### Genotyping

Mice were genotyped by PCR amplification of tail DNA with 4 sets of primers specific either for the genomic regions that were deleted in mutants but present in WT (5′-GTCTTGGTCTCATTCTAGGACACC and 5′-AACATGAGAGCTTCCAGTCCTCTC for *Per1* gene; 5′-AGTAGGTCGTCTTCTTTATGCCCC and 5′-CTCTGCTTTCAACTCCTGTGTCTG for *Per2* gene), or for the recombinant alleles present in mutants only (5′-ACAAACTCACAGAGCCCATCC and 5′-ACTTCCATTTGTCACGTCCTGCAC for *Per1^−/−^*, 5′-TTTGTTCTGTGAGCTCCTGAACGC and 5′-ACTTCCATTTGTCACGTCCTGCAC for *Per2^Brdm1^*).

### Immunohistochemistry

Eye globes were immersion-fixed in 4% paraformaldehyde in phosphate-buffered saline (PBS) overnight at 4°C. Eyeballs were rinsed in PBS, cut into two hemispheres and cryoprotected upon transfer to an ascending series of sucrose solutions (10%, 20% and 30% each for 1h) and then embedded (Tissue-Tek OCT compound; Thermo-Shandon, Pittsburg, PA, USA). Cryostat sections (10 μm thick) were permeabilized for 5 min with 0.1% Triton X-100 and saturated with PBS containing 0.1% bovine serum albumin, 0.1% Tween-20 and 0.1% sodium azide for 30 min. Sections were incubated overnight at 4°C with monoclonal anti-rhodopsin antibody Rho-4D2 [24]. Secondary antibody incubation was performed at room temperature for 2h with Alexa 488 anti-mouse IgG-conjugated antibodies (Molecular Probes Inc., Eugene, OR, USA). Cell nuclei were stained with DAPI (Molecular Probes). Slides were washed thoroughly, mounted in PBS/glycerol (1:1), and observed by an epifluorescence microscope (Nikon Optiphot 2). The number of phagosomes was quantified, as described previously by us [19]. Transverse sections (n=4/animal) were obtained from the central retina, covering the whole width of the retina from one periphery to the other. Taking the POS/RPE interface as a baseline, any immunopositive inclusion exceeding 1 μm lying within the RPE subcellular space was scored as a phagosome. Phagosomes were counted by aligning a 150 × 150 μm^2^ grid parallel with the RPE layer and displacing it dorsally and ventrally with respect to the optic nerve, along the POS/RPE interface, from the posterior to the superior margin. The phagosome counts are expressed as the sum of all 4 sections/eye.

### RT-qPCR gene expression analysis

Retinas were sampled immediately after sacrifice. A small incision was performed on the cornea with a sterile blade, lens and vitreous were discarded, and the retina was directly collected with sterile forceps and immediately frozen in liquid nitrogen and stored at -80 °C.

Retinas were homogenized in the RNable (Eurobio Scientific, France) solution by using a 23-gauge sterile needle and 1 ml syringe and mRNA extracted according to the manufacturer’s recommendations. Resuspended RNA was treated with DNAse | (0.1 U/μl, 30 min, 37°C - Fermentas) followed by phenol/chloroform/isoamylalcohol extraction and sodium acetate/isopropanol precipitation. RNA concentration and purity were measured using NanoDrop ND-1000V 3.5 Spectrophotometer (NanoDrop Technologies, Wilmington, DE, USA; A260/A280 and A260/A230 values were between 1.8 and 2). RNA quality was evaluated with the Bioanalyzer 2100 (Agilent Technologies; RNA integrity numbers were between 7.8 and 9).

500 ng of total RNA were reverse transcribed by using random primers and the “High Capacity RNA-to-cDNA” kit (Applied Biosystems, Foster City, CA, USA) following the manufacturer’s instructions. qPCR was performed using the 7300 Real-Time PCR System (Applied Biosystems) and the hydrolyzed probe-based TaqMan chemistry, with optimized Gene Expression Assays designed for specific mRNA amplification (**Table S1**). We used the TaqMan Universal PCR Master Mix with No AMPErase UNG (Applied Biosystems) and 1μl of cDNA in a total volume of 20 μl. The PCR program was; 10 min at 95°C and then 40 cycles of 15 s at 95°C and 1 min at 60°C. The fluorescence acquisition was performed at the end of the elongation step (7300 System Sequence Detection Software V 1.3.1 - Applied Biosystems). Each PCR reaction was done in duplicate. A dilution curve of the pool of all cDNA samples from one series was used to determine working dilution and to calculate the amplification efficiency for each assay (values were between 1.8 and 2 for all assays). No-template control reactions were performed as negative controls for each assay. One 96-well plate corresponded to the analysis of one gene. Data analysis was performed with qBase software (free v1.3.5) [25] and transcript levels were normalized using *Hprt* and *Tbp* that showed constant expression in their mRNA during the 24-h cycle (data not shown). Average gene expression levels within one experiment (one genotype) were set to 1, so that amplitudes (representing the maximal deviation from this 100% mean) could be compared between groups as was previously performed by Hiragaki and colleagues [26].

### Laser Capture Microdissection

Eye globes were enucleated under dim red light (<5 lux), embedded in OCT, snap-frozen and stored at -80°C until use. Eyes were cryosectioned at 10 μm thickness. Each eye provided 116 – 258 sections. All sections were dehydrated with ethanol and stained with Cresyl Violet staining (LCM Staining Kit, Ambion) and air-dried before microdissection with a Laser Microdissection System (LCM; PALM, Bernried, Germany). The RPE and photoreceptors were isolated with LCM (**Fig**. **S1**). The number of eye sections used for LCM RPE and photoreceptor isolation between genotypes was similar with 183 ± 10.18 (mean ± SD) slices used from WT eyes, whereas 201.3 ± 8.91 slices were used from double mutants (*P* = 0.19, Student’s *t*-test).

### RNA isolation for RNA sequencing

Total RNA was isolated using an RNeasy Micro kit (Qiagen Benelux, Venlo, The Netherlands), quantified with a Nanodrop (Isogen Life Science B.V., The Netherlands) and the quality was checked on a Bioanalyzer (Agilent Technologies, Amstelveen, The Netherlands). Sample RNA integrity (RIN) values for photoreceptors ranged from 7 to 9.8, except for 3 samples (RIN = 3.2, 4.1, 4.1). For RPE samples, RIN values ranged from 5 – 9.5.

### Library preparation and RNA sequencing

We used the KAPA mRNA HyperPrep kit (Illumina Platforms). For generating libraries, we used one batch of 20 ng of total photoreceptor (n = 8) RNA and 30 ng for the other three batches (n = 24) according to the manufacturer’s protocol (Illumina Platforms). For generating libraries from RPE samples we used 20 ng of RNA. RPE samples with low RNA yield were pooled. RPE libraries were generated in three batches.

The presence of cDNA was confirmed using flash gels (cat No. 57032, Lonza, Rockland, ME, USA). Libraries were 50 bp single-end sequenced using the Illumina HiSeq 4000 platform.

### Bioinformatics

The photoreceptor and RPE RNA-seq data were analyzed separately, but with the same software versions and parameter settings unless indicated otherwise. Raw sequencing data were subjected to quality control using FastQC (v.0.11.15), Picard Tools, and dupRadar [27]. All samples were of sufficient quality. Reads were trimmed for adapter sequences using Trimmomatic (v0.32) [28]. Trimmed reads were aligned to the mouse genome (Ensembl GRCm38.p6) using HISAT2 (v2.1.0) [29]. Gene level counts were obtained using HTSeq (v0.11) [30] with default parameters except --stranded=reverse and the mouse GTF from Ensembl (release 93). Statistical analyses were performed using the edgeR [31] and limma R (v3.5.0)/Bioconductor (v3.7) packages [32]. Genes with more than 2 counts in 4 or more samples (photoreceptors) or in 3 or more samples (RPE) were retained. Count data were transformed to log2-counts per million (logCPM), normalized by applying the trimmed mean of M-values method and precision weighted using voom [33]. Pairwise differential expression between the conditions of interest was assessed using an empirical Bayes moderated t-test within limma’s linear model framework, including the precision weights estimated by voom. Both for WT and *Per1^−/−^Per2^Brdm1^* a moderated F-test was used to determine which genes are differentially expressed between time-points. Resulting p-values were corrected for multiple testing using the Benjamini-Hochberg false discovery rate (FDR). An adjusted p-value < 0.05 was considered significant for photoreceptors. For the RPE an adjusted p-value of < 0.1 was considered significant. Additional gene annotation was retrieved from Ensembl (photoreceptors: release 94, RPE: release 98) using the biomaRt R/Bioconductor package. Gene ontology and pathway enrichment analysis was performed using g:Profiler [34]. We set all identified transcripts in our RNA-seq dataset as a reference background. We set an adjusted *P* < 0.05 as a threshold for significantly enriched pathways using the g:SCS method to correct for multiple testing [34]. We investigated interactions between protein products of the list of potential POS phagocytosis candidate genes by STRING analysis [35]. The 57 candidate genes encode for 49 proteins represented as nodes in the STRING network analysis. By setting the threshold to 0.25, we found 32 edges in the STRING network. Non-interacting nodes were not shown.

### Statistics

Data are represented as means ± SEM. Plots were generated using GraphPad Prism (La Jolla, CA, USA), SigmaPlot (Systat Software, San Jose, CA, USA) or R (Bell Labs, Murray Hill, NJ, USA). Normality of distribution was tested using the Shapiro-Wilk test. In case of non-normal distribution, the analysis was performed using ANOVA on ranks. Circadian expression profiles were determined using non-linear regression fitting to the equation y = y0 + c ∙ cos [2π (t-φ)/24], where y0 represents mesor, c amplitude and ϕ acrophase [36,37]. The function featured the following constraints: φ < 24, φ > 0 and c > 0. Gene expression profiles were considered to be rhythmic when significant fitting (*P* < 0.05) was observed to the equation y = y0 + c ∙ cos [2π (t-φ)/24]. Further analyses, where indicated, were performed using 1-way or 2-way ANOVA analysis followed by Holm-Sidak’s post hoc tests.

## Results

### Peak of rod outer segment phagocytosis is blunted in the retinas of *Per1^−/−^ Per2^Brdm1^* mice

The phagocytosis of photoreceptor outer segments is a highly rhythmic process occurring in a daily peak. This process persists in constant darkness, suggesting that it is driven by the circadian clock [12,38]. We tested the hypothesis that intact clockwork is required to sustain a rhythm of POS phagocytosis in constant darkness (DD). To that end, we used the *Per1^−/−^ Per2^Brdm1^* clock mutant mice which are behaviorally arrhythmic in DD [21].

We used age-matched (2 months old) wild-type and *Per1^−/−^ Per2^Brdm1^* mice, harvested eye globes at 8 time points over 24 h and analyzed anti-rhodopsin-stained phagosomes in the RPE (**Fig. 1a**). We quantified POS phagosomes at various time-points under DD conditions (n= 3 animals per genotype and per time point). A 2-way ANOVA analysis showed that the number of POS phagosomes was affected by genotype (WT versus *Per1^−/−^Per2^Brdm1^*, *P* < 0.001), time (*P* < 0.001) and an interaction between genotype and time (*P* < 0.001). Post-hoc analysis showed that phagocytic activity was rhythmic in wild type mice only, with 3-4 times more phagosomes at time-point CT1 compared with baseline (*P* < 0.001 for all time point comparisons) (**Fig**. **1b, c**; also confirmed by 1-way ANOVA, *F*7,16 = 34.49; *P* < 0.001). In contrast, in *Per1^−/−^ Per2^Brdm1^* mice, there was no obvious peak (1-way ANOVA, *F*7,16 = 2.35; *P* = 0.075). These results suggest that *Per1* and/or *Per2* is required for rhythmic POS phagocytosis.

**Figure 1.**
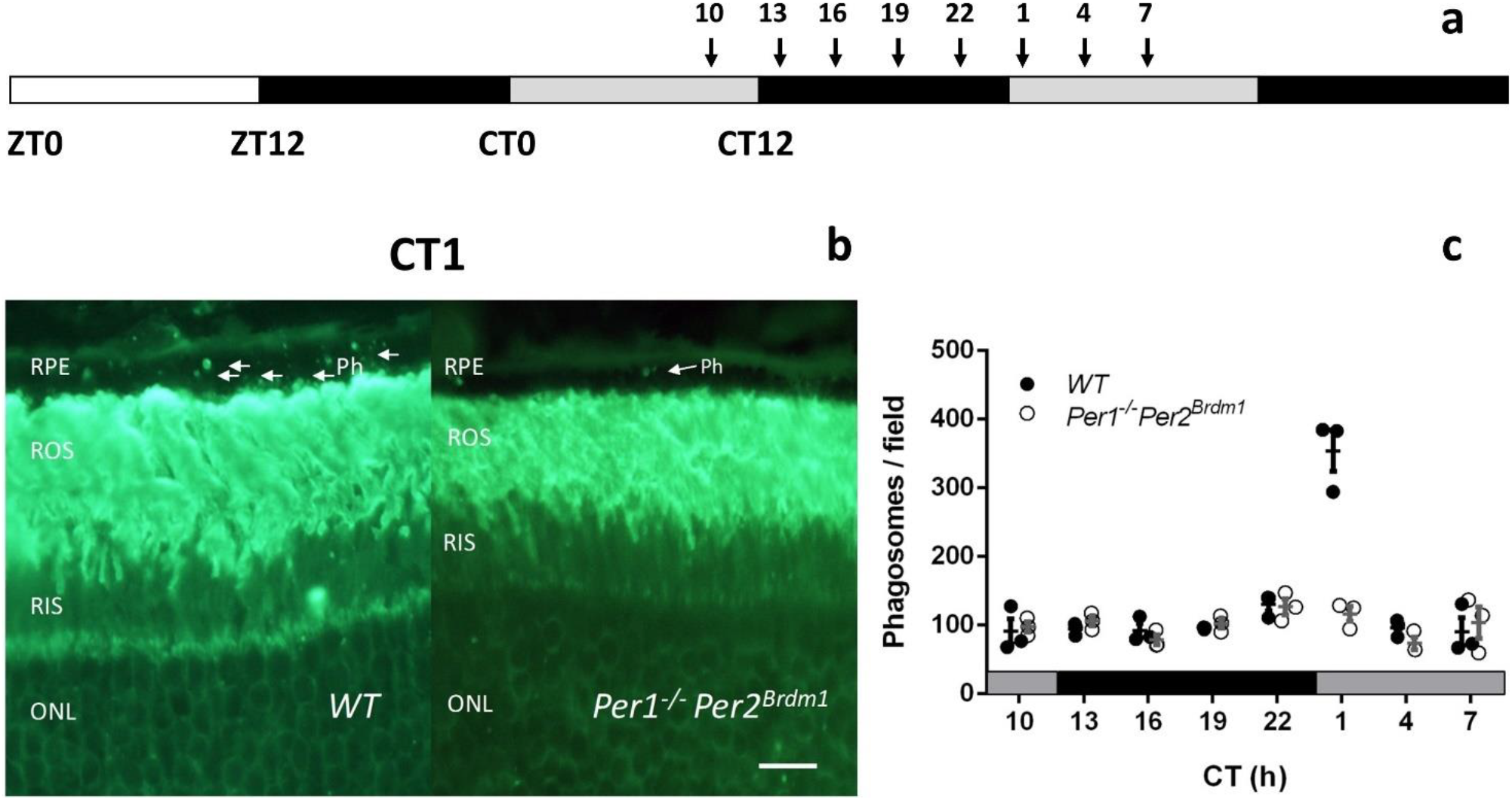
Mice lacking *Per1* and *Per2* show an impaired peak in POS phagocytosis. (a) WT and *Per1^−/−^ Per2^Brdm1^* mice maintained under 12h light (white bar) - dark (black bar) conditions were placed under constant darkness (DD, grey - black bars) and sacrificed at time-points indicated by arrows. (b) Representative image of Rho-4D2 stained phagosomes of WT and *Per1^−/−^ Per2^Brdm1^* retinas obtained at CT1 during the peak in phagocytosis in DD conditions. RPE – retinal pigment epithelium, ROS – rod outer segments, RIS – rod inner segments, ONL – outer nuclear layer, Ph – phagosomes. The scale bar is 10 μm. (c) Quantification of phagosomes in WT and *Per1^−/−^ Per2^Brdm1^* retinas under DD showed that *Per1^−/−^ Per2^Brdm1^* mice had no detectable peak in ROS phagocytosis. N = 3 / genotype / time-point. Graphs show mean ± SEM and values from individual samples are shown as dots.

### Molecular makeup of the retinal clock in absence of *Per1* and *Per2*

Since the peak of phagocytosis is attenuated in the mutant mice in DD, we hypothesized that the molecular clockwork is impaired in *Per1^−/−^ Per2^Brdm1^* retinas. To test this hypothesis, we sampled retinas from WT and *Per1^−/−^ Per2^Brdm1^* mice every 4h over 24h under DD, and quantified relative mRNA levels of clock genes by qPCR (**Fig. 2a**). Rhythmicity in expression profiles was assessed by cosinor analysis. These changes over the 24h cycle were mainly confirmed by 1-way ANOVA analysis (**Table S2**). Under DD conditions we found rhythmic clock gene expression for *Bmal1*, *Per1*, *Per2*, *Rev-Erbα* and *Rorb* in WT whole retinas (**Fig. 2b**, **Table S2**). Unexpectedly, in DD conditions, we found that in *Per1^−/−^ Per2^Brdm1^* mouse retinas also five clock genes were rhythmic: *Bmal1*, *Per3*, *Cry1*, *Cry2* and *Rev-Erbα.* Therefore, in contrast to our hypothesis, these results suggest *Per1* and *Per2* mutations do not significantly impair the rhythmicity of whole retinas in mice.

**Figure 2.**
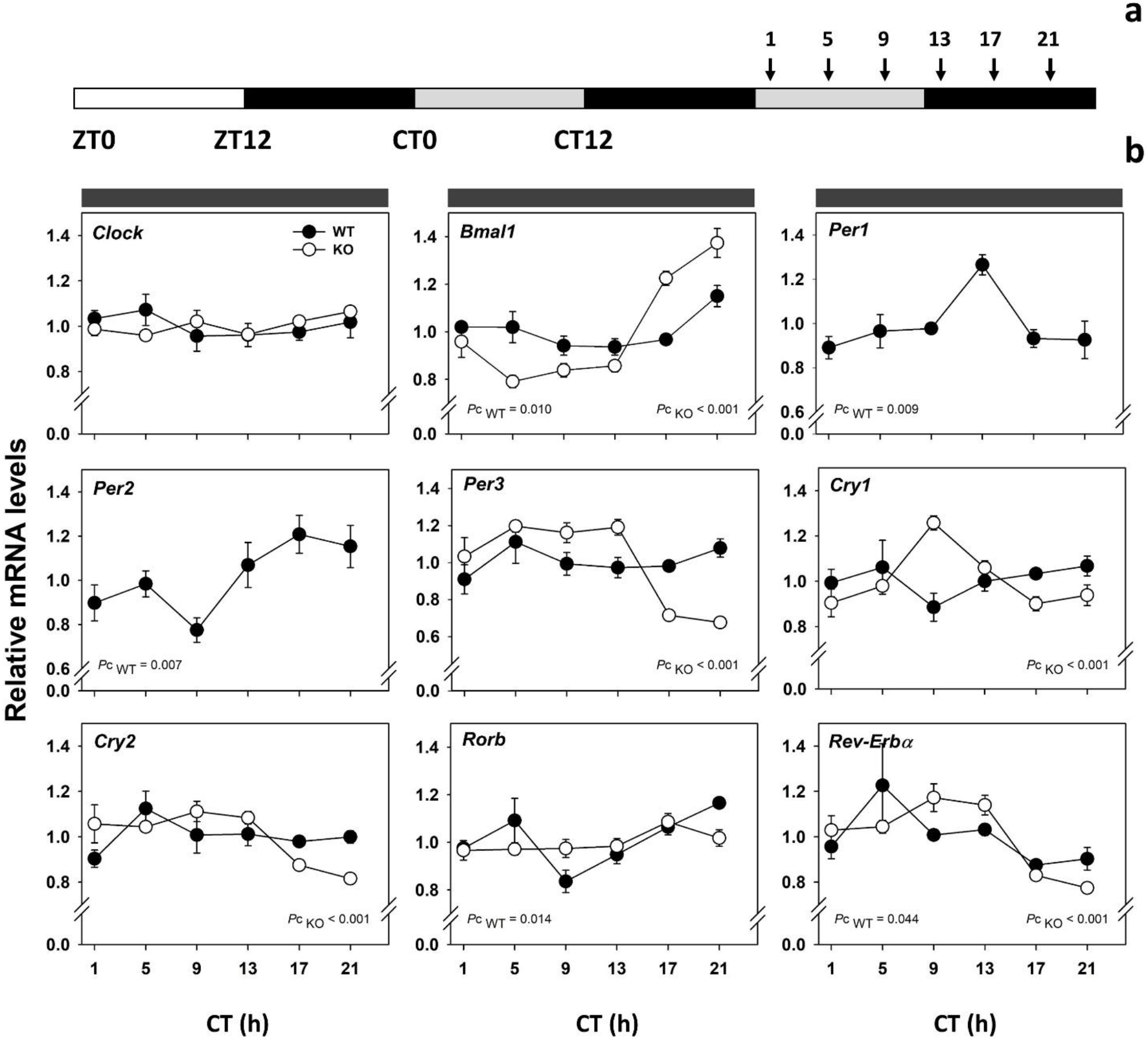
Clock gene expression profiles in WT (black dots) and *Per1^−/−^ Per2^Brdm1^* (white dots; KO) whole retinas under DD conditions. (a) Mice were placed under DD conditions, sacrificed at time-points indicated by arrows and their whole retinas were harvested. (b) QPCR analysis revealed that rhythmic gene expression was observed for *Bmal1*, *Per1*, *Per2*, *Rev-Erbα* and *Rorb* in WT retinas. Rhythmic expression was found for *Bmal1*, *Per3, Cry1, Cry2* and *Rev-Erbα* in *Per1^−/−^ Per2^Brdm1^* retinas. Values represent mean ± SEM. Significant temporal variations are indicated (*P* < 0.05). *Pc* – *P*-value of cosinor non-linear regression fitting to the equation y = y0 + c ∙ cos [2π (t-φ)/24], with y0 – mesor, c – amplitude and φ – acrophase. N = 3-4 for WT and 4-5 for double mutants / time-point.

### Transcriptomics analysis of WT mouse RPE and photoreceptors

To characterize the potential link between the circadian clock and the peak in POS phagocytosis, we first sought to characterize the time-affected transcriptomes of the RPE and photoreceptors. We harvested WT and *Per1^−/−^ Per2^Brdm1^* mouse eyes kept in DD at 4 time-points (CT19, 22, 1 and 10) (**Fig. 3a**). We laser-capture-microdissected the RPE and photoreceptors from each mouse eye (n = 4 / genotype / time-point), extracted RNA and performed RNA sequencing. In the RPE and photoreceptors, respectively, a total of 24 382 and 22 694 genes had sufficiently large counts to be retained in the statistical analysis. Next, we performed a pair-wise comparison of WT RPE and photoreceptor transcriptomes between consecutive time-points (**Fig. 3b** and **c**). In WT RPE, we found a large number of differentially expressed genes in comparisons between the expected peak in phagocytosis time-point CT1 and adjacent time-points (CT22 and CT10, respectively, **Fig. 3b**). In WT photoreceptors, most genes are differentially expressed in comparisons between CT10 and adjacent time points (CT1 and CT19, respectively, **Fig. 3c**). By contrast, in all pair-wise comparisons we found that only 3 genes differed significantly between time-points (i.e. were up-regulated at CT10 vs 19) in *Per1^−/−^ Per2^Brdm1^* RPE. We found that 1 gene was down-regulated at CT19 vs CT10 in *Per1^−/−^ Per2^Brdm1^* photoreceptors. Thus, these results suggest that the transcriptional program for initiating POS phagocytosis is likely in the RPE and not photoreceptors.

**Figure 3.**
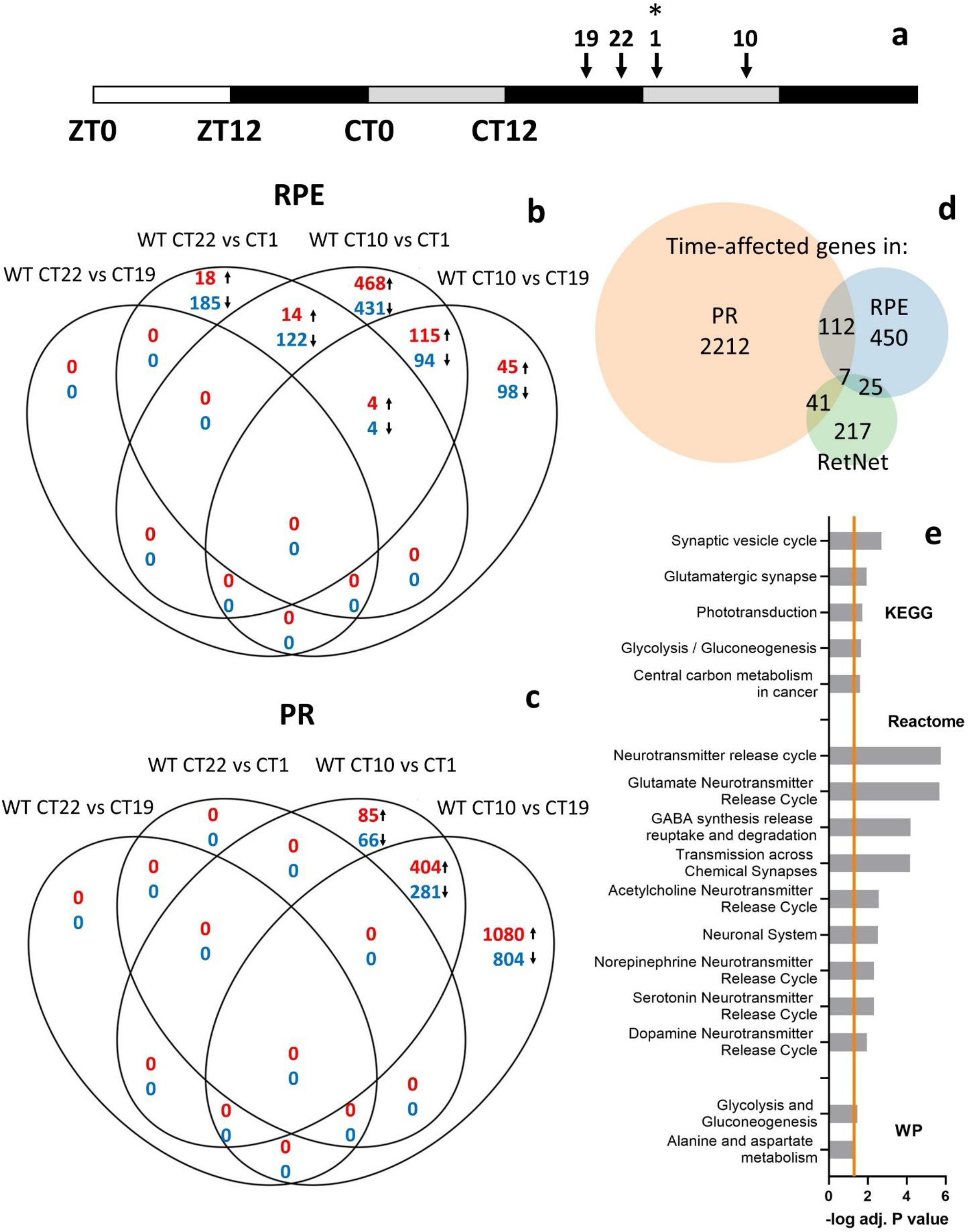
Transcriptional profiling of WT mouse RPE and photoreceptors. (a) Eyes were obtained under DD conditions from 4 successive time-points: before (CT19, CT22), during (CT1) and after (CT10) the expected peak in POS phagocytosis (n = 4 / time-point). RPE and photoreceptors were meticulously laser-capture-microdissected from each mouse eye, RNA was extracted and the transcriptomes were determined using RNA-sequencing. (b) In the RPE, a substantial number of genes were differentially expressed at time-points adjacent to the peak in POS phagocytosis – CT1. (c) By contrast, in photoreceptors (PR) most differential gene expression occurs around CT10. Red numbers represent the number of up-regulated differentially expressed genes, whereas blue ones are down-regulated. (d) A substantial number of identified transcripts showed a time effect in WT PR and RPE. There is considerable overlap (n=119) between time affected genes in these two tissues, a number of which overlap with the RetNet list of eye disease-related genes [39]. (e) Functional annotation (performed using g:Profiler) revealed that overlapping time-affected genes in RPE and PR are enriched in glucose metabolism and neurotransmission-related pathways. The orange line represents the significance level cut-off (adjusted *P* < 0.05). WikiPathways, Reactome and KEGG are databases of biological pathways.

Our differential expression analysis showed that 594 genes in WT RPE (=2.44% of all genes retained in the analysis) and 2 372 genes in WT photoreceptors (=10.45% of retained genes) varied over time-points (**Fig. 3d, Table S3, S4**). Among them are components of the circadian clock network (**Table S3, S4)**. Pathway analysis of time-affected genes in WT mice RPE revealed that, in addition to circadian pathways, phototransduction and metabolic-related pathways were functionally enriched (**Table S5**). Time-affected WT photoreceptor genes were enriched in circadian, metabolic, neurotransmission and DNA repair-related pathways (**Table S6**). Interestingly, 119 time-affected genes overlap in RPE and photoreceptors (**Table S7**), and are functionally enriched in glucose metabolism and neurotransmitter release-related pathways (**Fig. 3e**, **Table S8**). We also found that, respectively, 32 and 48 time-affected genes in the RPE and photoreceptors overlap with the RetNet list of eye disease-related genes [39] (**Fig. 3d, Table S9**). Thus, our results show that in the RPE and photoreceptors, a large number of genes and pathways vary in a time-dependent manner, a number of which are implicated in eye diseases.

### Potential molecular pathway that initiates POS phagocytosis

Our results suggested that the transcriptional events in the RPE might initiate POS phagocytosis (**Fig. 3b**). Our results also suggested that *Per1* and/or *Per2* are necessary for driving the peak in POS phagocytosis under DD (**Fig. 1**), but the molecular link is unclear. To characterize this link, we performed pair-wise comparisons between WT and *Per1^−/−^ Per2^Brdm1^* RPE transcriptomes. We found a substantial number of genes that were differentially expressed in *Per1^−/−^ Per2^Brdm1^* RPE compared to WT ones at the peak POS phagocytosis time-point CT1 (**Fig. 4a**). Next, we defined selection criteria for genes that potentially initiate POS phagocytosis (**Fig. 4b**). Considering that the peak in POS phagocytosis is lacking in *Per1^−/−^ Per2^Brdm1^* mice, we assumed that the genes that initiate POS phagocytosis are down-regulated in double mutant RPE compared to WT ones at CT1. POS phagocytosis occurs as a peak in WT mice on a molecular and functional level [40,41]. Thus, we selected genes that are both up-regulated at CT1 vs CT22 and down-regulated at CT10 vs CT1 in WT RPE. We removed possible photoreceptor “contaminants” from this list by using mouse signature cone and rod genes [42] and the Gene Ontology database (POS cellular component, GO:0001750). Using this strategy, we obtained a list of 57 candidate genes (**Fig. 4b, Table S10**). These genes are functionally enriched in neurotransmission related pathways (**Fig 4c, Table S11**). To reveal the interactions that protein products of these genes are involved in, we constructed a protein-protein interaction network using STRING [35] (**Fig. 4d**). Our list revealed a number of functional associations in which the protein products of candidate genes are involved in, most of which are associated with the term cell junction (highlighted in red in **Fig. 4d**). This cluster involves the interactions of *Syp*, *Gnaz*, *Pacsin1*, *Snap91*, *Camk2d* and *Camk2b* as identified in our STRING analysis. Thus, it is possible that POS phagocytosis might be initiated by the largest cluster identified in this analysis.

**Figure 4.**
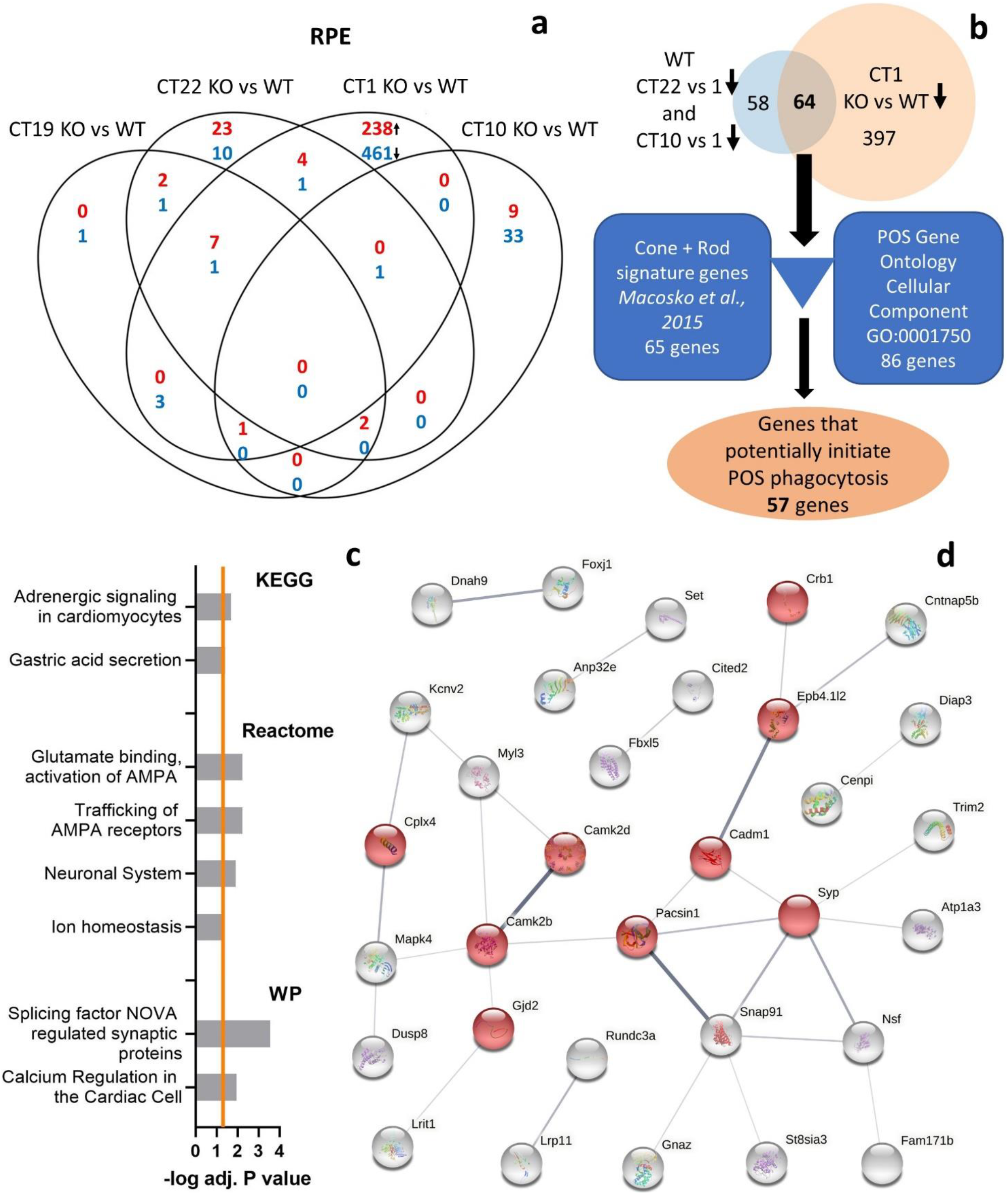
Identification of potential phagocytic pathways in RPE. (a) A comparison of WT and *Per1^−/−^ Per2^Brdm1^* (KO) RPE transcriptomes within each time-point revealed that most genes were differentially expressed during the peak phagocytosis time point - CT1. Red numbers represent the number of up-regulated differentially expressed genes, whereas blue ones are down-regulated. (b) Selection strategy for compiling the list of genes in the RPE possibly implicated in regulating POS phagocytosis. Signature rod and cone genes, [42] and the Gene Ontology term “Photoreceptor Outer Segment” were used to remove photoreceptor genes from the list of genes that potentially regulate POS phagocytosis. (c) Functional enrichment analysis using g:Profiler showed that these genes are enriched in neurotransmission related pathways from the WikiPathways (WP), Reactome, KEGG databases. The orange line represents the significance cut-off (adjusted *P* < 0.05). (d) STRING network analysis of protein functional associations of products of RPE genes implicated in initiating phagocytosis. Nodes represent protein products (n=57). Disconnected nodes are not shown. Edges represent protein functional associations. Interaction confidence scores range 0.25 - 0.99.

## Discussion

In the present study, we found no peak in POS phagocytosis in retinas of mice carrying a combined *Per1* and *Per2* mutation under constant darkness. Unexpectedly, gene expression analysis revealed that mutant retinas remained rhythmic under constant darkness, in contrast to mutant RPE and photoreceptors which showed no temporal variation. Using the purest possible RPE and photoreceptor sample material obtained by microdissection, we found significant differential gene expression in WT RPE at the peak phagocytosis time-point, but not in photoreceptors. Our results suggest a network of genes that potentially initiates POS phagocytosis in the RPE. These data challenge the view that molecular events in photoreceptors drive POS phagocytosis (via expression of phosphatidylserine “eat-me” signals) [18].

Retinal clocks are present in virtually all retinal layers [43–45,4] and are tightly coupled [4]. Coupling between retinal clocks contributes towards the precise timing of physiology within the retina [46]. In our study in *Per1^−/−^ Per2^Brdm1^* mice, constant darkness prevented any increased phagocytosis following subjective onset of day, a process known to be clock-regulated [12,47–50]. Thus, we speculated that constant darkness might impair the clockwork in *Per1^−/−^Per2^Brdm1^* whole retinas. The literature is not consistent regarding the effects of lighting conditions on clock gene expression in the whole retina. Studies either report no effects of DD on global retinal oscillations [45,51] or suggest that DD conditions dampen retinal rhythmicity [52,53,37]. Unexpectedly, our qPCR study revealed that clock gene expression remained rhythmic in both WT and *Per1^−/−^ Per2^Brdm1^* whole retinas. The origin of rhythmicity in mutant whole retinas is not known. It is most likely not due to input from the central clock, because retinal clocks are known to be independent from the SCN [3] and the SCN is considered arrhythmic based on locomotor activity of *Per1^−/−^ Per2^Brdm1^* mice in DD [21]. The source is most likely not in photoreceptors because in this study transcriptomics analysis of LCM-isolated *Per1^−/−^ Per2^Brdm1^* photoreceptors showed no temporal variations. Therefore, it is likely that rhythms in mutant whole retinas originate from retinal layers which display the most robust rhythms: e.g. the inner retina [4,43,44,51,37]. Considering that the number of oscillating genes differs considerably across mouse organs/tissues [54], it is possible that *Per1* and/or *Per2* mutations impact the RPE and photoreceptor clocks disproportionally more than the clockwork of other retinal cells. Regardless of the reasons, these results suggest that (global) retinal rhythmicity is not sufficient for driving the peak of POS phagocytosis.

The phagocytosis of POS is a rhythmic process occurring roughly one-hour after light onset [12,47–50]. This process is critical for retinal health as demonstrated by retinal degeneration displayed in both human patients [16] and animal models [55,15,56]. Some literature stresses the importance of precise timing of POS phagocytosis in maintaining retinal health [55,56]. This view is corroborated by our finding that a number of eye disease-related genes vary across time-points in the RPE and photoreceptors. However, it was recently reported that dopamine D2 receptor knockout mice had no peak in POS phagocytosis and displayed no apparent retinal pathologies [57]. Regardless, the molecular pathways responsible for driving this peak are not known [5,17,18]. By using immunohistochemistry and quantifying ingested POS in clock mutant mouse retinas we showed that *Per1* and/or *Per2* are necessary (molecular clock) components for the transient surge in POS phagocytosis.

The prevailing view is that POS phagocytosis is initiated by the externalization of phosphatidylserine “eat-me” signals on the POS membrane [17,18]. However, we found that microdissected WT photoreceptors did not differ in gene expression 3h or 6h before the peak in POS phagocytosis. By contrast, in WT RPE, we found that a number of genes were differentially expressed at the phagocytic peak time-point compared to the 3h earlier one. In addition, at the peak phagocytosis time-point, we found a vast number of differentially expressed genes in *Per1^−/−^ Per2^Brdm1^* RPE compared to WT ones. These results suggest that POS phagocytosis is initiated by the RPE. This possibility is indeed plausible because the RPE was shown to display sustained rhythms in various models: in vivo [57–61]; ex vivo [62–64] and in cell culture models [64–68]. Importantly, the phagocytic machinery is rhythmic in these cells [57,67,41,64]. Furthermore, in an arrhythmic BMAL1 knockout cell culture model there was no rhythm of phagocytic activity [67].

Finally, we proposed a network of genes for regulating ROS phagocytosis in the RPE. The candidate genes in this list are enriched in the ion homeostasis pathway. This is expected as previous studies implicated ion channels in POS phagocytosis such as voltage gated sodium channels [69] and the L-type calcium channel Cav1.3 [55]. The list also contains known genes implicated in POS phagocytosis such as *Mfge8* [70] and *Myl3* [71]. Cell junctions were also enriched in the candidate gene list, among which *Gjd2* encodes for a gap junction protein. It is possible that increased gap junction expression enhances the connectivity of the RPE at the peak phagocytic time-point. That might, in turn, lead to a synchronized and sharp phagocytic peak across the whole RPE. However, it should also be noted that a number of genes in the list have not been, sufficiently characterized e.g. *Gm13112*, *Gm13735*, *Gm16701*, etc. Therefore, our list of candidate genes provides ample opportunities for investigation for the research community.

The strength of this approach is the use of the purest possible sample material obtained from LCM. In addition, we considered the rhythmic nature of POS phagocytosis by using samples from multiple time points. We also compared our results with an arrhythmic mouse model that lacked this peak phagocytic activity. There are some limitations in our approach. For example, the genes implicated in initiating phagocytosis might not be down-regulated after the peak phagocytic time-point. It might be that at the peak phagocytic time-points, the down-regulated genes repress RPE phagocytic activity. It is also possible that genes in the list might be “contaminants” originating from POS fragments that are ingested by the RPE. Despite the imperfections, this list will be a valuable tool for studying the POS phagocytosis pathway.

In conclusion, our study reveals that *Per1* / *Per2* are necessary circadian clock components for driving the rhythm of POS phagocytosis. Our results show that *Per1* and *Per2* mutation does not impair the rhythmicity of the whole retina. Our data suggests that the molecular pathways that initiate POS phagocytosis are most likely initiated by the RPE by genes functionally enriched in neurotransmission related pathways.

## Abbreviations

Bmal1: Brain and Muscle ARNT-Like 1
BP: biological process
CPM: counts per million
Cry: Cryptochrome
DD: constant darkness
DEGs: differentially expressed genes
FDR: false discovery rate
Hprt: Hypoxanthine Phosphoribosyltransferase
KEGG: Kyoto encyclopedia of genes and genomes
LCM: laser capture microdissection
LD: light-dark cycle
MF: molecular function
Per: Period
POS: photoreceptor outer segment
ROS: rod outer segment
Ror: RAR-related orphan receptor
RPE: retinal pigment epithelium
SCN: suprachiasmatic nucleus
Tbp: TATA-Box binding Protein
WP: WikiPathways
WT: wild type
ZT: Zeitgeber time

## Acknowledgements

We thank Anneloor L.M.A. ten Asbroek and Nguyen-Vy Vo for technical assistance. We thank Dr. D. Sage, Dr. S. Reibel and N. Lethenet from the Chronobiotron (UMS 3415) for animal care and Dr. U. Albrecht (University of Freiburg) for the *Per1^−/−^Per2^Brdm1^* mice. This project has been funded with support from the NeuroTime Erasmus+ grant (European Union), Rotterdamse Stichting Blindenbelangen (Netherlands), Nelly Reef fund (Netherlands), Stichting voor Ooglijders (Netherlands), Stichting tot Verbetering van het Lot der Blinden (Netherlands) and Retina France (France).

