## Supplemental figure for "Core Circadian Clock Genes *Per1* and *Per2* regulate the Rhythm in Photoreceptor Outer Segment Phagocytosis"

***** Equal first author contribution

^ Equal last author contribution

### Supplementary figure

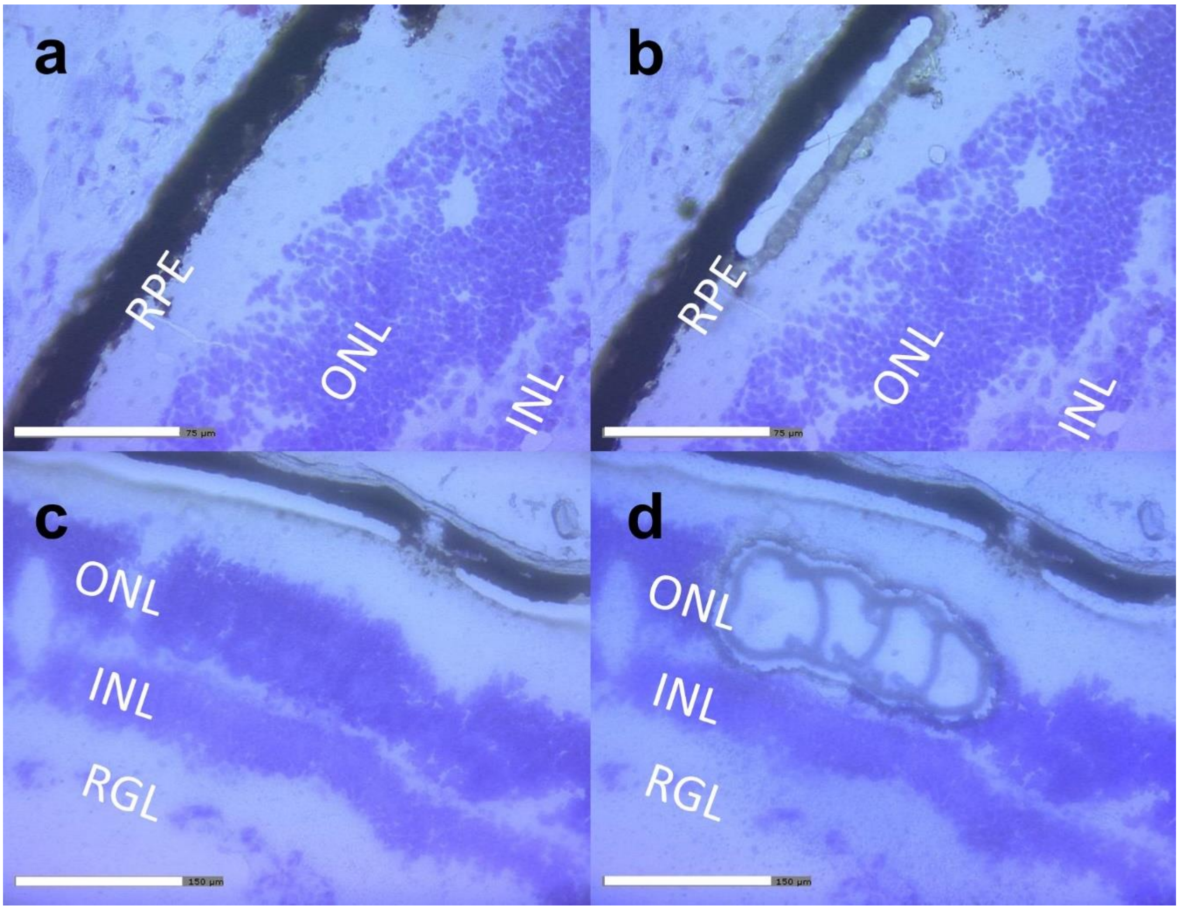


**Figure S1**. Laser capture microdissection of photoreceptors from WT and $Per1^{-/-}Per2^{Brdm1}$ mouse eyes. Eyes were cryosectioned at 10 µm thickness. Nuclei were stained using cresyl violet. (a) Micrographs of an eye section before and (b) after laser capture microdissection of the RPE are shown. The scale bar is 75 µm. (c) Micrographs are shown of slice before and (d) after microdissection of photoreceptors. The scale bar is 150 µm. ONL – outer nuclear layer; INL – inner nuclear layer and RGL – retinal ganglion layer.
